# Navigating the window: structural bottlenecks in avian craniofacial integration revealed by the ontogenetic patterning of shape variance

**DOI:** 10.64898/2026.09.17.752413

**Authors:** Diane Hu, Jay Devine, Wei Liu, Benedikt Hallgrímsson, Ralph S. Marcucio, Nathan M. Young

## Abstract

**Background:** Successful palatal morphogenesis requires the precise spatiotemporal choreography of independent facial prominences. Our previous macroevolutionary analyses identified a convergence in amniote facial growth at primary palatal fusion, associated with reduced shape variability. We hypothesize that this constraint operates within individual species, representing a shared structural bottleneck where brain growth acts as a critical morphogenetic co-factor. To test this idea, we analyzed high-resolution 3D micro-CT images of embryonic chickens (*Gallus gallus*, N=217) spanning primary palatogenesis. Utilizing 3D geometric morphometrics (3DGM), we quantified a continuous "developmental morphospace" integrating the face and brain.

**Results:** Facial shape follows a highly nonlinear developmental trajectory. Using a continuous sliding-window analysis, we demonstrate that local morphological disparity undergoes a significant system-wide collapse coinciding with primary palatal fusion. Partial Least Squares (PLS) confirms this morphospace pivot is tightly coordinated with neurocranial development. Post-fusion, the modules uncouple: facial allometry switches from convergent to divergent projecting growth, while the brain undergoes spatial encapsulation, subsequently flatlining its morphological disparity.

**Conclusions:** Craniofacial morphogenesis can be described in terms of a tightly regulated biphasic developmental hourglass. The physical tethering of facial prominences creates a brief structurally constrained "shape window" essential for successful fusion. Deviations from this bottleneck — whether driven by altered prominence trajectories, increased local variance, or disruptions to the underlying forebrain landscape — push embryos into unoccupied morphospace, providing a unified mechanical and predictive framework for the spatiotemporal origins of non-syndromic cleft lip.

## Introduction

Normal craniofacial development requires the spatiotemporal choreography of multiple, distinct tissues that must combine to produce a highly integrated structure critical to an organism’s survival (Chai & Maxson, 2006). Modeling this facial development presents numerous challenges. Thus, predicting with confidence exactly when, how, or why facial morphogenesis may go awry remains elusive (Dixon *et al*., 2011). Improving our quantitative ability to model these growth trajectories provides valuable information on how localized developmental processes contribute to severe pathologies such as cleft lip specifically and generalized facial dysmorphology more broadly (Wyszynski & Beaty, 1996). Such modeling is a crucial step toward translating descriptive insights from basic developmental science into actionable mechanisms for genetic counseling and targeted therapy.

Morphogenesis of the primary palate occurs early in embryonic facial development and involves the orchestration of paired prominences (the frontonasal, lateral, and maxillary processes) that must independently grow, contact, and fuse to generate a functional upper jaw. Given the sheer mechanical complexity of this process, minor deviations can propagate into severe dysmorphology, making cleft-lip one of the most common structural birth defects in humans, affecting approximately 1 in 700 live births globally (Jiang & Bush, 2006). Consequently, most studies investigating craniofacial anomalies have focused heavily on the role of prominence growth, frequently linking cleft-lip to intrinsic reductions in prominence size or altered proliferation rates (Wang *et al*., 1995; Diewert & Lozanoff, 2002). Size mismatches or delayed growth phases increase the mechanical risk that individual prominences will fail to successfully contact and integrate (Diewert *et al*., 1993; Young *et al*., 2007; Boughner *et al*., 2008; Parsons *et al*., 2011). This makes early, abnormal prominence growth a critical determinant of later palatogenesis failures (Trasler, 1968; Young *et al*., 2014).

Previously, we constructed a comparative developmental morphospace designed to characterize the macroevolutionary patterns of facial growth across deeply divergent amniotes, including mammals, reptiles, and avians (Young *et al*., 2014) [**Figure 1A: Comparative Developmental Funnel**]. Despite identifiable, species-level differences in early embryonic stages, and the diversity present among adults, the facial trajectories of highly divergent species converged as prominence growth proceeded toward fusion. Furthermore, empirical simulations within "unoccupied" regions of the developmental morphospace during this narrow period of minimal variance successfully predicted that aberrant prominence growth inevitably results in unsuccessful contact and CL, an outcome subsequently confirmed by experimental perturbation (Young *et al*., 2014). We hypothesized that shared structural constraints strictly limit the functional solutions for successful palatogenesis, resulting in this deeply conserved developmental shape convergence.

**Figure 1.**
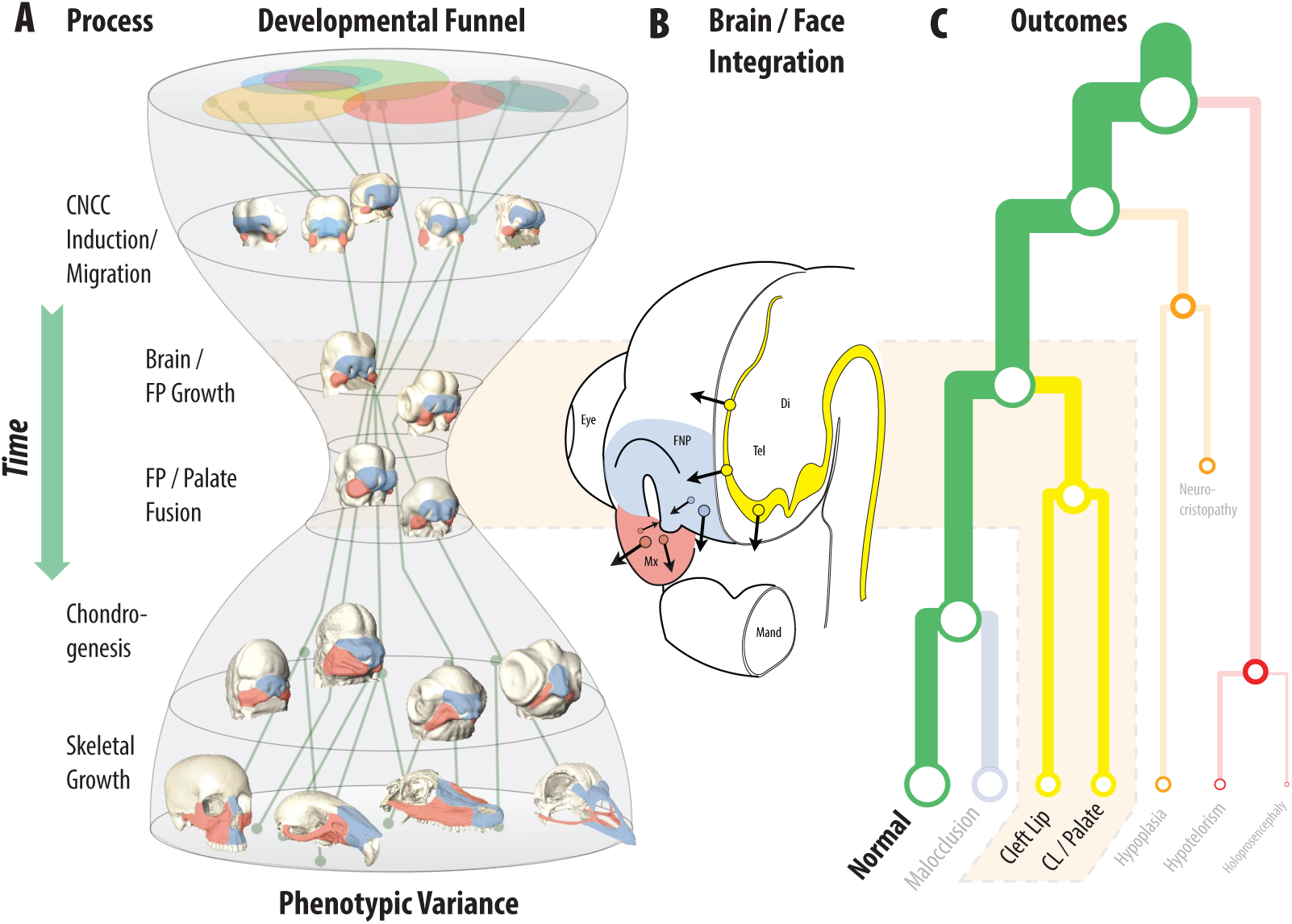
Conceptual Framework of Craniofacial Convergence and Structural Integration. **(A)** The Comparative Developmental Funnel: A macroevolutionary morphospace demonstrating that despite high phenotypic diversity in early embryos and adults across amniotes, facial shape results from a series of conserved processes that converge into a highly constrained shape window during the period of primary palatal fusion (Young *et al*., 2014). **(B)** Structural Adjacency of the Brain and Face Integration: Schematic illustrating the intimate physical and signaling relationship between the developing forebrain and the facial prominences (frontonasal: blue; maxillary: red; neuroepithelium: yellow). The brain acts both as a signaling center and as an expanding physical platform that dictates the spatial distances prominences must traverse to achieve contact and fusion (Marcucio *et al*., 2011). **(C)** Developmental Outcomes are Process Dependent: Disruption of these processes leads to hierarchical and contingent disruptions to facial outcomes, where earlier events may have syndromic effects that cascade into additional outcomes (Hallgrímsson *et al*., 2009). The face is particularly prone to disruptions in events associated with a fusion bottleneck that may lead to isolated cleft-lip outcomes.

If this hypothesis is true, similar, highly regulated patterns of restricted shape variability must necessarily operate *within* individual species during development. However, successful palatogenesis depends on more than just the intrinsic growth of the facial prominences. It relies heavily on both external molecular signals and the broader structural environment in which those prominences grow, most notably, the developing brain (Boughner *et al*., 2008; Parsons *et al*., 2011; Hu *et al*., 2015) [**Figure 1B: Brain/Face Structural Adjacency**]. The brain serves dual roles during embryogenesis: it acts as a critical signaling center (e.g., orchestrating SHH-signaling from the frontonasal ectodermal zone) and a massive structural platform, forming a highly integrated mechanical relationship with the developing face (Marcucio *et al*., 2011; Parsons *et al*., 2011; Chong *et al*., 2012). For example, as the developing forebrain increases in volume, its surface area rapidly expands. This expansion mechanically spreads the associated placodes and inherently alters the physical, three-dimensional distance that the facial prominences must traverse to achieve contact (Hu *et al*., 2015).

Therefore, we hypothesize that contact and fusion events must coincide with a tightly constrained, system-wide developmental window within individual species, and that brain shape and growth rate act as significant conditional variables determining the phenotypic landscape of the face. To rigorously test these predictions, we present high-resolution developmental morphospaces of both face and brain morphogenesis in a dense ontogenetic series of the chicken (*Gallus gallus*). By mapping the continuous localized morphological disparity of these tissues throughout development, our analyses reveal mutually distinct biphasic morphogenetic pathways that lead to facial dysmorphology, offering novel quantitative insights into the generalized structural mechanisms of clefting [**Figure 1C: Brain/Face Structural Adjacency**].

## Methods

### Embryo Collection and Imaging

To construct a continuous high-resolution dataset, we collected chicken (*Gallus gallus*) embryos (N=217) across a dense developmental window, sampling every 4 to 6 hours from Hamburger-Hamilton (HH) stages 16 to 30 (Hamburger & Hamilton, 1951). This corresponds roughly to 72–148 hours of post-fertilization incubation (N=∼10 embryos per HH stage). Embryos were carefully dissected in cold PBS, fixed in 4% paraformaldehyde (PFA), and treated with a 1% Lugol’s iodine contrast solution to enhance the radiopacity of soft tissues, allowing for the precise resolution of both external and internal structures. All specimens were subsequently imaged via high-resolution micro-computed tomography (μCT, Scanco VivaCT 40) at a resolution of 10 μm (Degenhardt *et al*., 2010; Wong *et al*., 2013).

### Image Registration and Segmentation

We utilized an automated, non-linear volumetric registration pipeline to generate stage-specific templates (i.e., mean volumes) for automated phenotyping (Percival *et al*., 2019; Devine *et al*., 2020) **[Figure 2: 3D Segmentations]**. This involved spatially aligning stage-specific μCT images to an iteratively defined average target image at increasingly higher resolutions using hierarchical affine and non-linear deformations via the ANIMAL algorithm (Devine *et al*., 2020). We then recovered and concatenated the transformations in preparation for propagating segmentation labels and landmarks from the template space to the original images. For every individual sample and stage-specific template, we generated two highly detailed 3D segmentations using the software *Amira* (v. 4.1.1, Mercury Systems): (1) an external surface isolating the developing “face,” and (2) an internal volume isolating the “brain” by approximating the dense underlying neuroepithelium.

**Figure 2.**
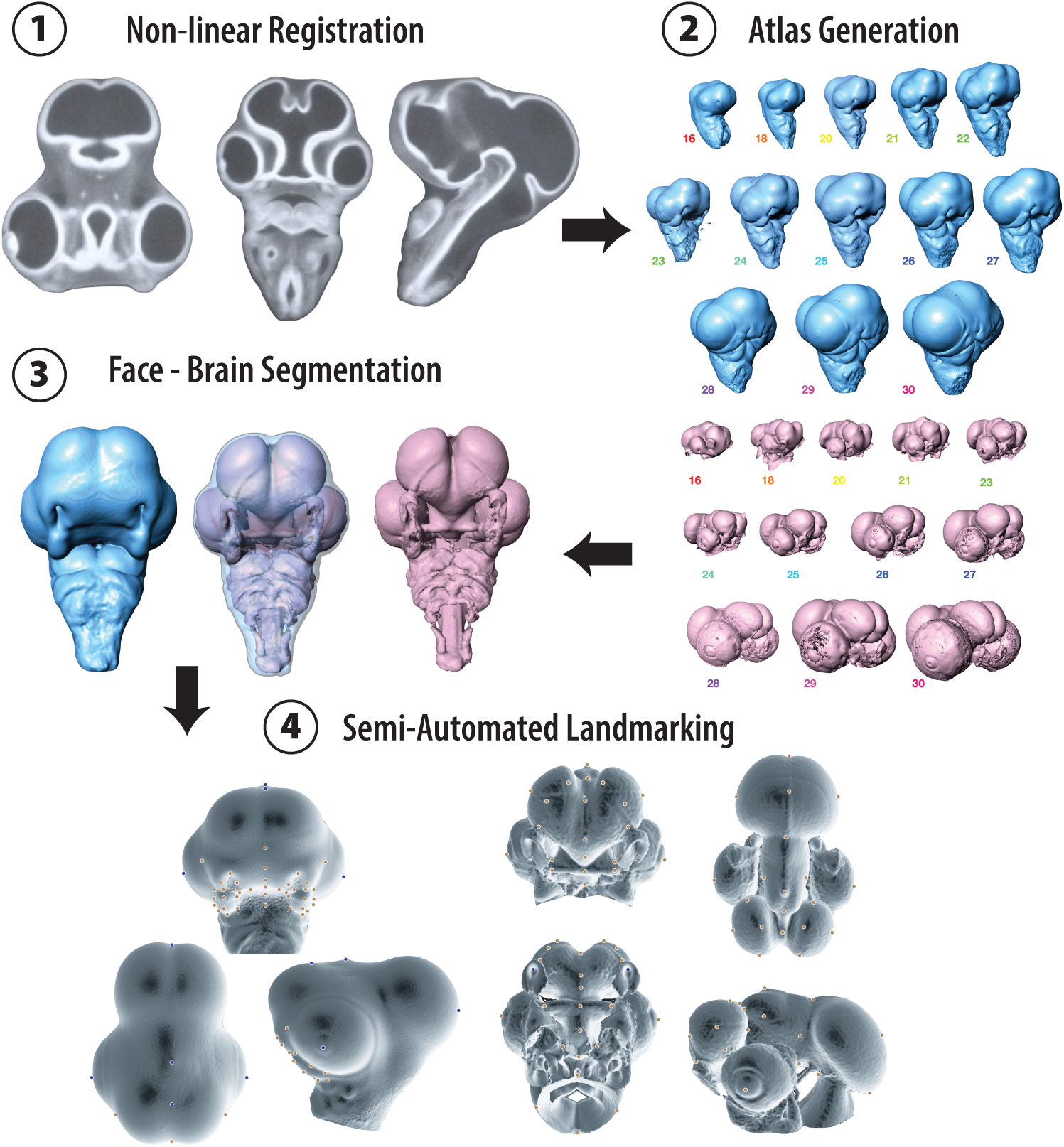
3D Morphometric Phenotyping of the Embryonic Chicken Head. High-resolution, quantitative modeling of face and brain morphogenesis from Stages 16 to 30. (1) Embryos were imaged via micro-CT with iodine contrast then (2) subjected to non-linear registration to create stage atlases. (3) 3D segmentation of both the external facial ectoderm (Face) and the internal dense neuroepithelium (Brain) was performed for each individual embryo. (4) A reproducible atlas of 3D landmarks was applied to both face and brain segmentations (N=40, each) to capture size and shape variation across developmental time after Generalized Procrustes Analysis (GPA) (see **Supplemental** Figure 1).

### Landmarking & Geometric Morphometrics (3DGM)

A standard reproducible set of 3D anatomical landmarks was placed on the stage-specific face (N=40) and brain templates (N=40), serving as a unified morphological atlas [**Supplemental Figure S1: Landmarks**]. A semi-automated landmark estimation protocol (*Landmark Editor*, UC Davis) was then applied to rapidly identify homologous points across all individual embryos, followed by rigorous manual adjustment and verification (Devine *et al*., 2020). The raw 3D coordinate configurations were subjected to Generalized Procrustes Analysis (GPA) to mathematically isolate true shape variation by removing the confounding effects of scale, position, and spatial orientation (Zelditch *et al*., 2004; Adams & Otarola-Castillo, 2013).

### Developmental Morphospace & Partial Least Squares (PLS)

We quantified within-species developmental morphospaces for both the face and brain modules using Principal Components Analysis (PCA) of the Procrustes shape coordinates (Mitteroecker & Gunz, 2010). To explicitly test the hypothesis of structural interaction between the modules, we utilized a two-block Partial Least Squares (PLS) analysis. This technique assesses the significant patterns of covariation between the face and brain by decomposing the covariance matrices derived from a joint superimposition of both coordinate sets into a unified Procrustes shape space (Rohlf & Corti, 2000; Klingenberg, 2009). All analyses were performed in the software *MorphoJ* v.1.08.02 (Klingenberg, 2011).

### Continuous Shape Variance (Sliding Window)

Traditional developmental studies often calculate variance by binning specimens into discrete chronological stages (Zelditch *et al*., 2004). However, this approach is frequently confounded by developmental heterochrony or sampling effects, where embryos of identical chronological or incubation age may exhibit vastly different levels of morphological maturity. To quantify changes in morphological disparity as a continuous, high-resolution function of developmental time, we implemented a sliding-window analysis.

First, specimens were sorted strictly by their scores along Principal Component 1 (PC1), which served as an objective, continuous proxy for developmental age [**Supplemental Figure S2: Size / Shape / Time**]. Next, a sliding window of fixed sample size (N=15) was translated along the sorted PC1 axis with a step size of one specimen. Within each local window, a localized consensus shape was dynamically calculated. True morphological disparity (local Procrustes variance) was then computed for each window as the sum of squared Procrustes distances of the N specimens to their *local* consensus, divided by N-1. By dynamically recalculating the consensus within each step, this method isolates the shape variance orthogonal to the primary directional growth trajectory, yielding a mathematically rigorous continuous measure of morphospace constraint and expansion.

## Results

### Developmental Morphospace: Face

PCA ordination of facial shape indicates that the first two principal axes are sufficient to capture the predominant patterns of developmental change, accounting for 84.4% of the total variance **[Figure 3A-B: Face Morphospace]**. Embryonic age distributes smoothly and predictably across PC1, isolating the monotonic passage of developmental time. Multivariate regression of shape on centroid size (64.4% variation, p<0.0001; *r*_v_=5.3°, p<0.0001), hours of development (67.6% variation, p<0.0001; *r*_v_=4.2°, p<0.0001), HH stage (65.5% variation, p<0.0001; *r*_v_=6.1°, p<0.0001), all indicate similar outcomes and shape trends, although a smaller proportion of total variation **[Supplemental Figure S2: Size / Shape / Time]**. These results confirm PC1 is a superior proxy for biological age and growth, capturing the progressive anterior and medial extension of the maxillary prominence and the accompanying antero-ventral movement of the frontonasal prominence **[Supplemental Figure S3: Face PC warps]**.

**Figure 3.**
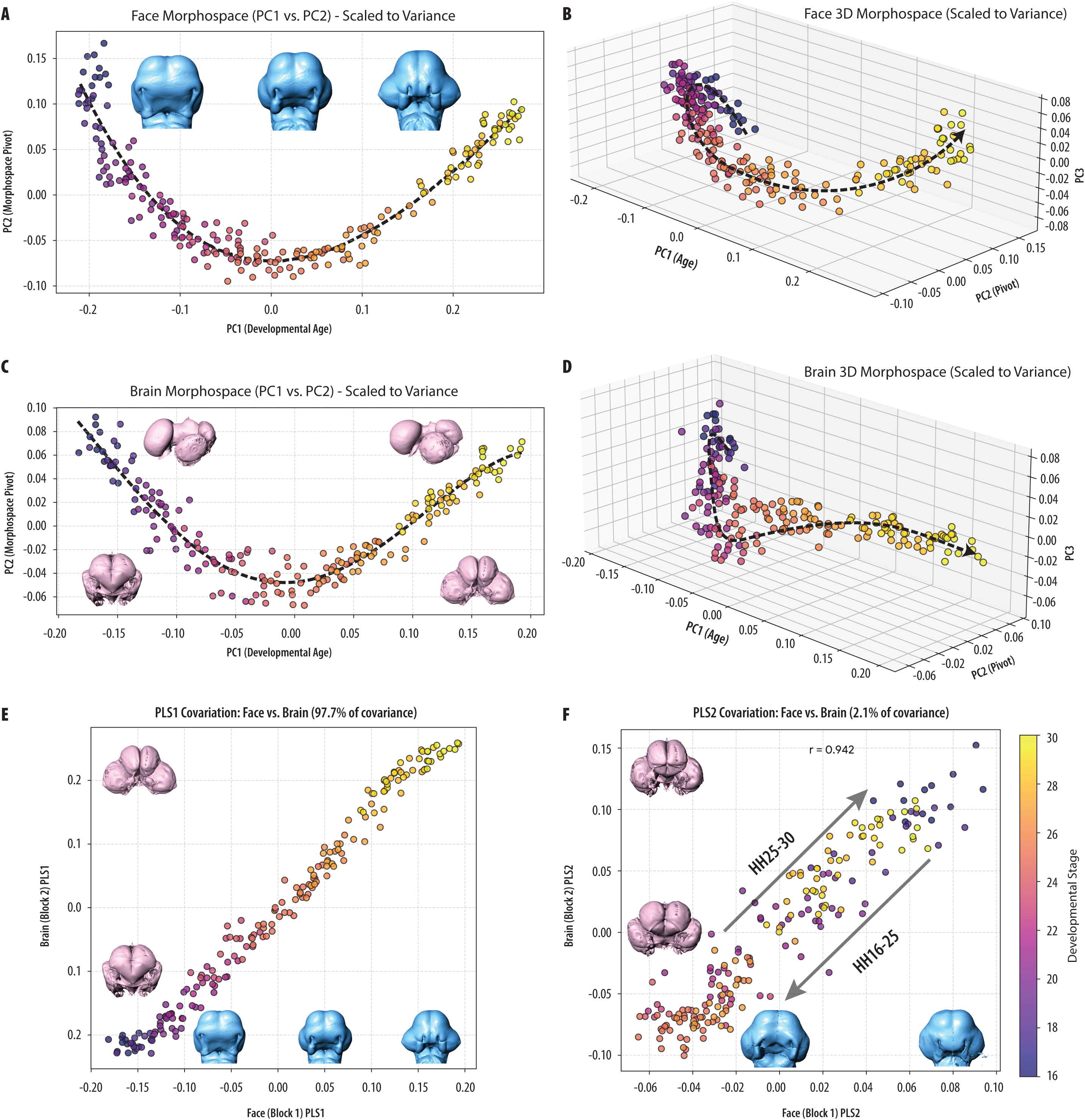
Biphasic Developmental Morphospaces and System-Wide Integration. (A-B) Principal Components Analysis (PCA) of facial shape. PC1 acts as a continuous proxy for developmental age, while PC2 captures a dramatic non-linear allometric switch ("U-turn") corresponding to the fusion of the facial prominences. 3D plots illustrate how embryonic shape traverses morphospace in a highly nonlinear fashion, with PC3 capturing anteroventral extension of the FNP. **(C-D)** PCA of internal brain shape reveals a mirrored trajectory reversal at the exact same developmental window, driven by the anterior rotation of the eyes and shifting forebrain proportions. **(E-F)** Two-block Partial Least Squares (PLS) analysis confirms strong system-wide covariation. PLS1 captures coordinated baseline growth (97.7% covariation, *r*=0.991), while PLS2 isolates the synchronized morphospace pivot (*r*=0.942), demonstrating that the face and brain navigate the fusion bottleneck as a highly integrated unit.

However, evaluating the secondary axis of variation, PC2, reveals a dramatic non-linear allometric switch in the morphospace characterized by proportional differences in the maxillary and frontonasal prominences. Initially, the maxillaries are small relative to the frontonasal, but subsequent growth drives a highly directional convergence of maxillary landmarks toward fusion with the frontonasal. Immediately after fusion, the trajectory of growth becomes abruptly divergent: the anterior extension of the maxillary prominences slows, while the frontonasal prominence continues to project significantly forward, switching their relative proportions again. This transition executes a sharp U-turn or "hook" in the morphospace, shifting the facial tissues from a convergent to a divergent proportional pattern occurring roughly at stage HH25. The superiority of a quartic fit (r^2^=0.91) to the data over a quadratic fit (r^2^=0.89) confirms that this inflection reflects a highly regulated biologically determined ontogenetic trajectory, rather than a product of a random walk or morphological drift (Bookstein, 2013; Polly & Motz, 2016).

### Continuous Facial Disparity (The "Double-Dip" Hourglass)

Tracking the local Procrustes variance of the face via the continuous sliding-window analysis reveals a dynamic multi-phasic model of developmental constraint (Scharloo, 1991; Green *et al*., 2017) **[Figure 4A: Continuous Face Variance]**. During early development (pre-fusion), spatial disparity is exceptionally high (variance ≈0.0070) as the independent prominences grow with a high degree of structural freedom. However, as the trajectory approaches the fusion window (Stages 24–25), morphological variance collapses by nearly 50%. This drop represents the primary structural bottleneck, as the once-individuated soft-tissue prominences mechanically tether together during fusion into a continuous integrated upper jaw structure.

**Figure 4.**
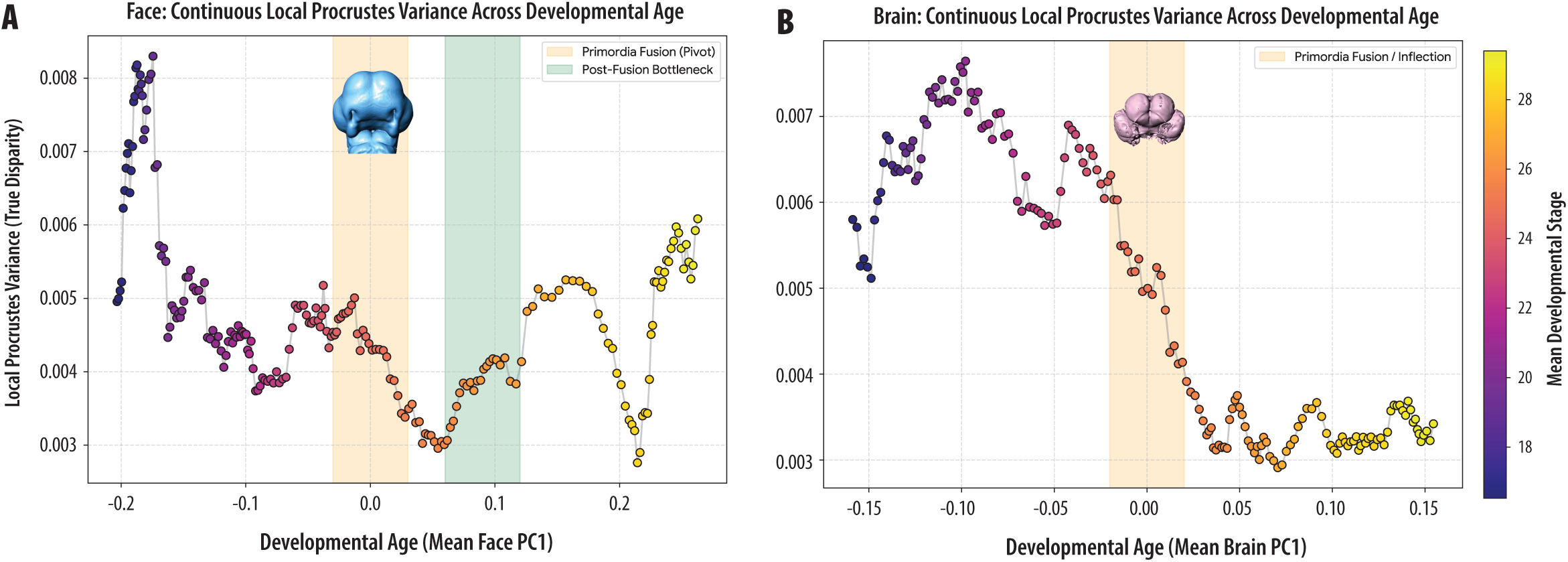
Continuous Morphological Disparity Reveals Post-Fusion Uncoupling. Sliding-window analysis of local Procrustes variance across continuous developmental time (PC1). **(A)** The face exhibits a "double-dip" hourglass trajectory: an initial massive collapse in variance during soft-tissue tethering (Fusion Bottleneck, orange), followed by a secondary constraint during prenasal chondrogenesis (green). Post-cartilage formation, facial disparity increasing as projecting allometry resumes in the elaboration of the nascent beak. **(B)** The brain exhibits a fundamentally divergent response. Following the shared variance collapse at the fusion bottleneck, brain disparity does not recover, reflecting its physical encapsulation and strict spatial constraint within the rigid neurocranium.

Following this fusion bottleneck, facial variance rebounds slightly before hitting a second drop (variance ≈0.0027) around Stages 28–29. While the absolute magnitude of this secondary dip varies across specific sampling windows [**Supplemental Figure S4: Disparity Robustness**], its consistent presence corresponds to the onset of prenasal chondrogenesis. Biologically, this may be explained as a temporary locking-in of the facial geometry by a newly forming rigid cartilaginous scaffold. Once this foundation is fully established by Stage 30, the mechanical constraint lifts. Facial disparity begins to expand outward once again (variance >0.0060) as secondary projecting traits of the now fully integrated facial prominences begin to re-accumulate significant phenotypic variance, such as species-specific beak allometry.

### Developmental Morphospace: Brain

The ordination of the internal brain shape shares a striking highly coordinated resemblance to the facial morphospace, with 73.4% of variation captured on PC1 and PC2 **[Figure 3C-D: Brain Morphospace]**. PC1 tracks ordered stage-specific neurocranial changes, most notably the division of the telencephalon and the massive anterior-posterior rotation of the developing eyes (centroid size: 63.4% variation, p<0.0001; *r*_v_=1.2°, p<0.0001; hours of development: 61.5% variation, p<0.0001; *r*_v_=2.3°, p<0.0001; HH stage: 60.3% variation, p<0.0001; *r*_v_=2.2°, p<0.0001). Like the face, the brain’s PC2 axis isolates a strong non-linear trajectory reversal that pivots through the morphospace at the exact same developmental window (∼HH25). Around Stages 23–25, the relative proportions of the cranial architecture undergo a shift: the rapid width expansion of the brain slows significantly, while the eyes rotate anteriorly **[Supplemental Figure 5: Brain PC warps]**. This internal reorganization gives the distinct impression of mechanically "squeezing" the anteriorly extending facial prominences from behind (Hu *et al*., 2015).

Crucially, the continuous sliding-window variance for the brain demonstrates a fundamentally divergent response to this shared fusion bottleneck **[Figure 4B: Continuous Brain Variance]**. Like the face, the brain exhibits a high degree of initial morphometric disparity that collapses sharply during the Stage 24–26 fusion window. However, unlike the "double-dip" and subsequent expansion seen in the face, the brain does not increase its variance again in the period sampled. Instead, from Stage 26 through Stage 30, the morphological disparity of the brain remains at the constrained minimum (variance ≈0.0031).

### System-Wide Integration (Face-Brain PLS)

A two-block PLS analysis confirms that the face and brain remain highly integrated throughout this morphological pivot (Rohlf & Corti, 2000; Klingenberg, 2009) [Figure 3E-F: PLS Trajectory]. PLS1 captures the magnitude of their shared baseline ontogenetic growth (97.7% of total covariation, r=0.991). More importantly, PLS2 (2.1% covariation, r=0.942) isolates the observed synchronized trajectory reversal. This strong correlation confirms that the allometric switch in the facial prominences and the structural spatial shifting of the forebrain and eyes occur not as isolated local events, but as a tightly coupled system-wide morphogenetic response.

## Discussion

### The Morphospace Pivot and System-Wide Integration

Our analysis of the embryonic chicken head reveals a highly integrated biphasic developmental trajectory defined by coordinated allometric switches. By employing both discrete stage-based ordinations and continuous sliding-window analyses, we capture a morphospace "pivot" accompanied by a collapse in spatial variance midway through the observed developmental window. This structural bottleneck coincides with the physical fusion of the facial primordia, functioning as a system-wide morphogenetic inflection point. Prior to fusion, the facial prominences develop as unconstrained modules, charting a highly variable directional vector through shape space. Upon fusion, these previously autonomous tissues tether into a unified structural ring, redirecting the geometry of craniofacial expansion and creating a coordinated shift in the morphospace.

Crucially, the two-block PLS analysis confirms that this pivot is not localized to the external face. The internal neuroanatomy executes an identical synchronized trajectory reversal. This intense degree of covariation demonstrates that the convergence of the external face primordia acts as a system-wide spatial constraint, either directly pulling the developing brain through the same developmental bottleneck via mechanical tethering or arising cooperatively because of the brain’s intimate structural adjacency.

### Divergent Disparity: Encapsulation vs. Projection

While the brain and face are highly integrated as they pass through the fusion event, calculating their continuous morphological disparity reveals that these tissue modules handle post-fusion physical constraints in fundamentally different ways reflecting their structural roles. The face follows a dynamic "double-dip" hourglass pattern. An initial drop in variance driven by soft-tissue fusion is followed by an apparent secondary constraint phase likely dictated by prenasal chondrogenesis (Stages 28–29). Once this underlying cartilaginous scaffold is permanently established, the face essentially becomes a projecting module. It is physically free to expand outward into open space, allowing species-specific phenotypic disparity to rapidly accumulate and expand.

In contrast, the brain functions as an encapsulated module. While it shares the initial variance collapse during fusion, its disparity does not increase again as it becomes physically confined within the forming rigid neurocranial capsule. This structural uncoupling mirrors evolutionary diversity: the extreme highly variable evolutionary plasticity of the amniote face exists in direct mechanical contrast to the strictly regulated and more highly conserved internal architecture of the neurocranium.

### Mechanisms of Clefting: Trajectories and the Phenotypic Landscape

Previously, we argued that successful primary palatogenesis depends entirely on the embryo’s ability to successfully navigate a narrow highly restricted “shape window” during this crucial developmental phase. The continuous variance results presented here strongly support this hypothesis, and further delineate how specific spatial disruptions to this structural integration lead mechanistically to cleft lip through distinct trajectory deviations:

1. **Increased Variability:** Elevated morphological variance in prominence shape rapidly expands the confidence interval of phenotypic outcomes [**Figure 5A: Clefting Mechanisms**]. This effectively pushes a significantly greater proportion of individual embryos into unoccupied cleft-lip-liable morphospace, resulting in pathology even if the mean developmental trajectory of the population remains entirely unaffected (Young *et al*., 2007; Green *et al*., 2019).
2. **Trajectory Shifts:** Altering the relative growth ratio between specific prominences (e.g., favoring frontonasal growth over maxillary growth) fundamentally alters the available contact area and restricts the available fusion time [**Figure 5B: Clefting Mechanisms**]. Previous experimental work selectively modulating *Shh* activity demonstrates that altering these delicate proportions actively steers embryos directly into unoccupied morphospace, virtually guaranteeing a cleft-lip outcome (Young *et al*., 2010, 2014; Chong *et al*., 2012; Li *et al*., 2013).
3. **The Brain as a Phenotypic Landscape:** Because the facial prominences are structurally tethered to the anterior neural tube, the forebrain acts as the physical spatiotemporal landscape across which facial morphogenesis must occur (Marcucio *et al*., 2011). If intrinsic brain growth is abnormally accelerated, the temporal window available for facial contact is dramatically shortened, and the midface is mechanically forced to spread apart before coalescence can occur [**Figure 5C: Clefting Mechanisms**]. Conversely, if brain growth is delayed, the embryo faces the risk of premature spatially abnormal fusion events (Chong *et al*., 2012).

**Figure 5.**
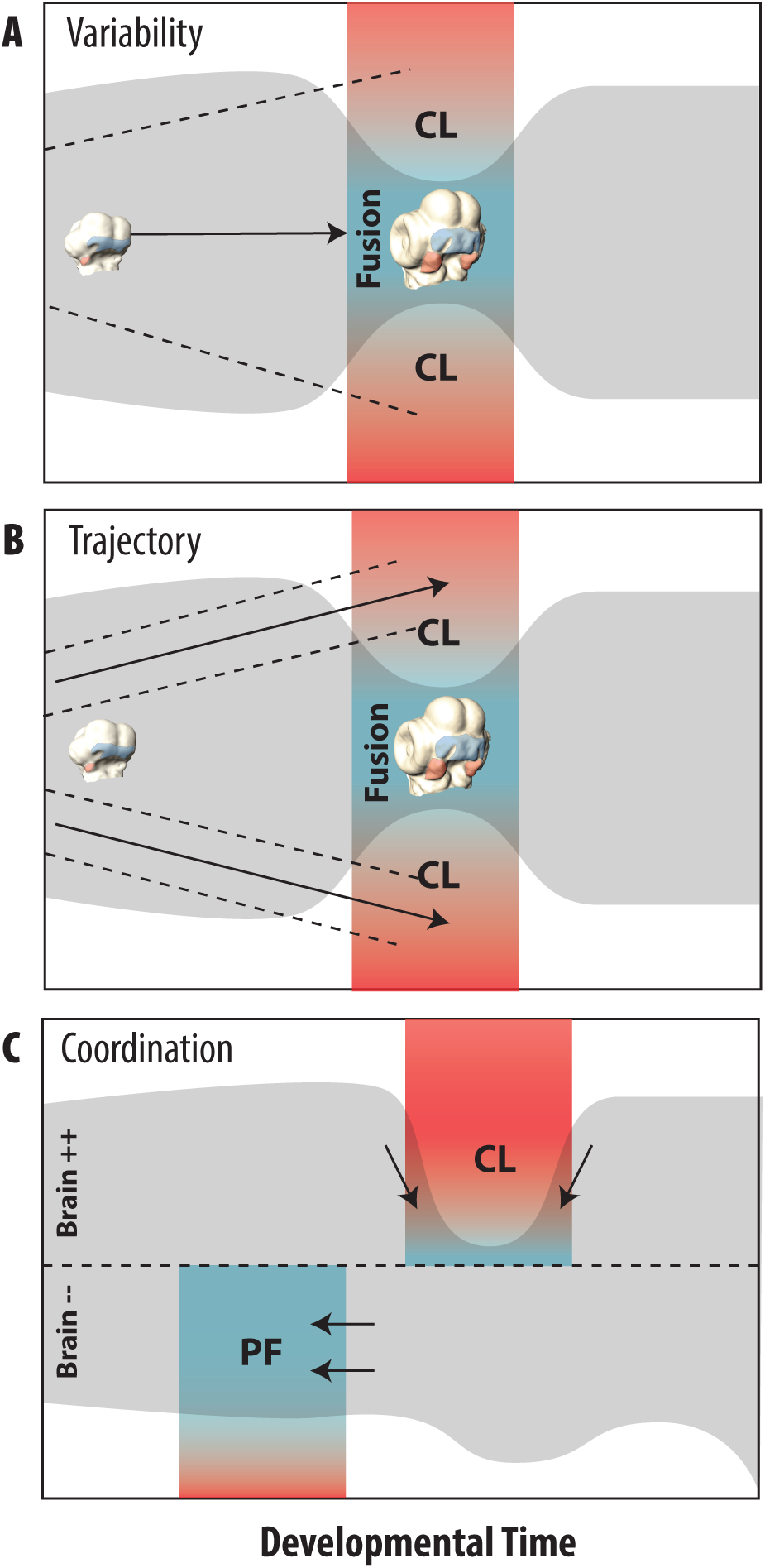
Mechanistic Origins of Non-Syndromic Cleft Lip. Conceptual models demonstrating how specific spatiotemporal deviations around the structurally constrained "shape window" lead to cleft lip (CL) pathologies. **(1) Increased Variance:** Elevated morphological disparity in growth trajectory pushes the tails of the population distribution outside the required bounds for fusion, resulting in cleft-lip even if the mean trajectory is normal. **(2) Shifted Trajectory:** Altered growth ratios between prominences cause the mean developmental trajectory to physically miss the required spatial target for fusion. **(3) Altered Forebrain Landscape:** Disruptions to brain growth rate act as a physical wedge; accelerated forebrain growth widens the landscape, creating an unbridgeable spatial gap (CL), whereas decelerated growth narrows the landscape, risking premature fusion.

### Evolutionary Implications and Conclusion

The morphospace pivot and concurrent variance collapse observed in avian ontogeny are highly likely to be conserved across disparate amniote lineages, functioning as a fundamental developmental hourglass where deeply conserved geometric boundaries must override lineage-specific variation (Boughner *et al*., 2008; Young *et al*., 2014). Framing pathological facial clefting as a simple spatiotemporal mismatch during this universal bottleneck allows us to extend macroevolutionary observations to successfully explain non-syndromic facial variation within a single species.

In summary, chicken craniofacial development is governed by a tightly integrated biphasic trajectory anchored by a severe structural bottleneck at Stages 24–25. The fusion of the facial prominences briefly synchronizes the face and brain into a narrow “shape window” that is necessary for successful palatogenesis. Post-fusion, the tissue modules fundamentally decouple: the neurocranium permanently encapsulates the brain to restrict its future disparity, while the face navigates prenasal chondrogenesis before entering a highly variable, unconstrained phase of projecting beak growth. This structural uncoupling illustrates how a tightly regulated developmental hourglass gives way to extensive morphological plasticity, providing a unified mechanical explanation for both macroevolutionary craniofacial diversity and the origins of non-syndromic facial clefting.

## Acknowledgements

This research was funded in part by grants from the National Institutes of Health and National Institute of Dental and Craniofacial Research: F32DE018596 (NY), R56DE029124 (NY, RM, BH), R01DE018234 (RM, BH), R01DE019638 (RM, BH).

## SUPPLEMENTAL FIGURE LEGENDS

Figure S3. **Face PCA Warps.**

Comparison of facial surface warps for PC1-3 relative to the mean shape shown in various orientations. Warped face surfaces were generated by applying the associated eigenvector of each PC to the mean shape scaled to the maximum (+) or minimum (-) value of the associated PC scores represented in the data.

Figure S4. **Comparison of Continuous Local Variance Metrics.**

**A-B**. We varied the window size as a low-pass filter on the variance data. Results confirm that the fundamental biological pattern we observe is extremely robust to methodological choices. Regardless of whether you look at 10 embryos at a time or 25, the macro-patterns are identical. The face consistently shows a "double-dip" (early fusion bottleneck and later chondrogenesis bottleneck) followed by expansion. The brain consistently shows an initial fusion bottleneck followed by later variance constraint. **C-D**. Dual-axis chart for the Face and Brain dataset utilizing an alternative fixed Δ*PC*1 window (Δ = 0.05). **The Purple Line (Left Axis)** is the true morphological disparity (local Procrustes variance), calculated strictly over consistent intervals of developmental time rather than an arbitrary number of embryos. **The Teal Stepped Fill (Right Axis)** tracks *N*, the actual number of embryos that happened to fall within each sliding 0.05 PC1 slice. By holding developmental time constant, we reduce any potential artifacts caused by sample clustering while revealing consistent and robust results.

Figure S5. **Brain PCA Warps.**

Comparison of brain surface warps for PC1-3 relative to the mean shape shown in various orientations. Warped brain surfaces were generated by applying the associated eigenvector of each PC to the mean shape scaled to the maximum (+) or minimum (-) value of the associated PC scores represented in the data.

---

**Figure S1.**
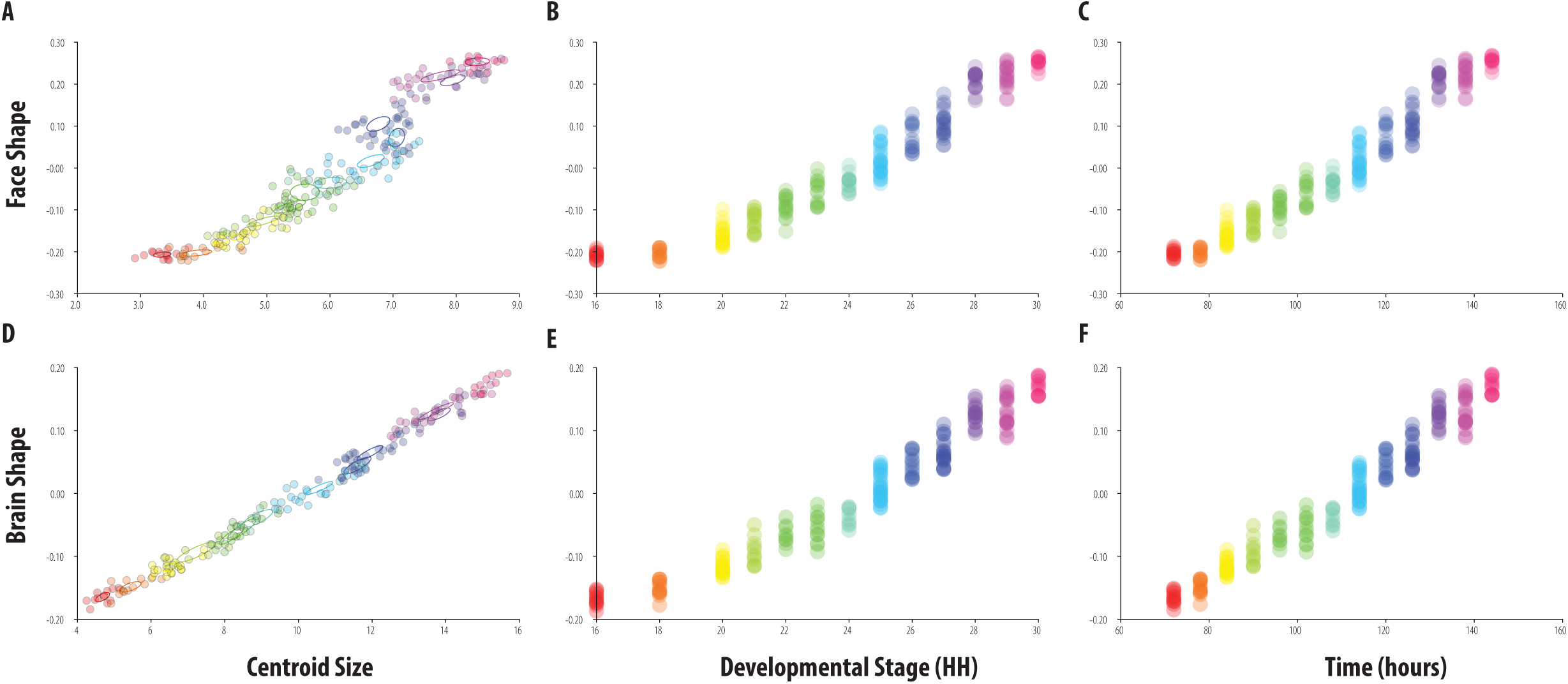
Landmarks. Face **(A)** and brain **(B)** landmarks shown in various orientations. Blue indicates the landmark was included in the associated shape dataset. Yellow indicates the landmark was used as a reference for atlas generation and to improve semi-automated registration prior to manual adjustment but was not included in data analysis.

**Figure S2.**
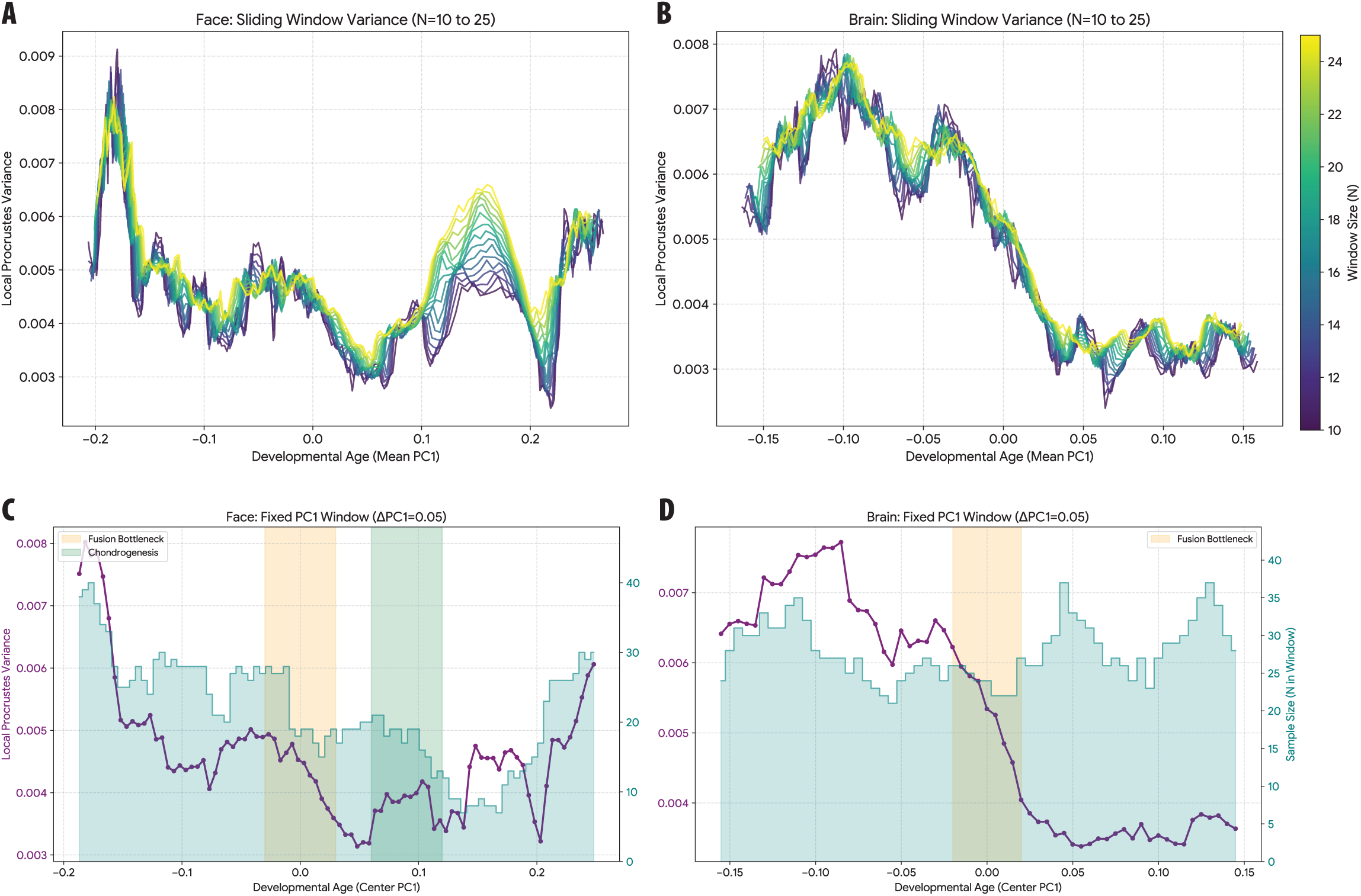
Comparison of Shape Size Metrics. Multivariate regression of shape of the face **(A-C)** and the brain **(D-F)** on centroid size **(A,D)**, developmental stage **(B,E)**, and incubation time **(C,F)**. In all cases there is significant relationship, consistent with a developmental trajectory, and all largely indistinct from that estimated from PC1.

